# Identification of spatial compartments in tissue from *in situ* sequencing data

**DOI:** 10.1101/765842

**Authors:** Gabriele Partel, Markus M. Hilscher, Giorgia Milli, Leslie Solorzano, Anna H. Klemm, Mats Nilsson, Carolina Wählby

## Abstract

Spatial organization of tissue characterizes biological function, and spatially resolved gene expression has the power to reveal variations of features with high resolution. Here, we propose a novel graph-based in situ sequencing decoding approach that improves recall, enabling precise spatial gene expression analysis. We apply our method on in situ sequencing data from mouse brain sections, identify spatial compartments that correspond with known brain regions, and relate them with tissue morphology.

## INTRODUCTION

Highly multiplexed spatial expression analysis of genes is essential to uncover the spatial organization of biological processes in relation to different regions of tissues and organs. Spatial atlases for well-studied organs like the brain exists^1^, but matching individual samples to a reference atlas is challenging as 2D sections may be angled and not well-aligned, as well as shape and size of organs differ between individuals. Furthermore, specimens from human organs are often sampled from a part of an organ, and it is generally not trivial to identify the exact original location and orientation of the analyzed sample. Some regions, such as the pyramidal cell layer of CA1 hippocampus in the brain, can be well defined based on morphology, whereas other regions are more difficult to identify as the morphological variations are small when cellular labeling is limited to nuclear stainings or a finite set of fluorescent markers. Instead, spatial compartments in tissue can be mapped by taking advantage of the spatial organization of gene expression from known biomarkers within each tissue sample.

Several methods for measuring gene expression while preserving spatial information have been developed over the past years. Generally described, there are two approaches to preserve spatial information: One approach is to either imprint a spatial reference on a grid or to carefully record spatial location prior to collection and sequencing of single-cell RNA^2,3,4,5,6^. The other approach is the parallel profiling of large numbers of mRNAs using barcodes decoded directly in the tissue sample^7,8,9,10^. All approaches come with benefits and drawbacks, such as spatial resolution, depth of sequencing, accuracy and throughput^11^.

Targeted in situ sequencing using padlock probes and localized rolling circle amplification^7^ provides submicron localization of dozens to thousands of RNA species simultaneously in cells and entire tissue sections, and recent advancements in automation^12^ have led to higher detection efficiency and shorter protocol times. Investigated genes are targeted with carefully designed barcoded padlock probes, locally amplified and sequenced by repeated fluorescent staining and imaging cycles. The resulting image data consists of five fluorescent channels for each sequencing cycle: One channel showing cell nuclei, and four channels with fluorescent signals representing the four bases of the genetic code (T, G, C, A). An additional general fluorescent marker detecting all amplified padlock probes is often added and used as a reference. The resulting fluorescent signals appear as bright spots in a noisy background caused by light scattering and autofluorescence in the tissue. Moreover, fluorescent spots may be subjected to slight spatial jitter between cycles, and often have a blurry appearance due to the limited signal-to-noise ratio of diffraction limited microscopes. Although super resolution methods may produce data with better signal-to-noise ratios, they would limit throughput at the same time. All these factors, together with increasing signal density at increasing multiplexity, make it challenging to distinguish real fluorescent signals from noise and background structures and consequently accurately decode barcode sequences.

Previous approaches to decode in situ sequencing data have relied on comparably simple morphological pre-processing, a global intensity threshold of the general stain image, followed by a watershed segmentation to resolve clusters, and a final decoding based on extracting fluorescence intensities per color channels and sequencing cycle^7^. Furthermore, a sequence quality metric based on relative signal intensities, was applied to filter away noise from true signals. Although providing promising results, many signals were missed in order to maintain a good precision-recall-ratio and consequently limiting the decoding resolution (i.e. number of transcripts per spatial unit).

## RESULTS

Here, we present a novel in situ sequence decoding pipeline that aims to be as inclusive as possible while maintaining high precision, thus pushing spatial gene expression profiling of tissue samples to large sample coverage, high multiplexity, and high decoding resolution, enabling unprecedented identification of tissue compartments. We detect local fluorescent maxima using a generous threshold in order to maximize sensitivity and delay the decision of which detected fluorescent signal candidates contribute to an expected sequence. Rather than trying to manually tune morphological filters to define true signals we adopt a Convolutional Neural Network (CNN). Briefly, we use a pre-trained CNN (Supplementary Fig. 11) (trained on a different dataset) and use extracted features as a probability prediction to describe how similar a signal candidate is compared to a true signal judged by visual examination. Thereafter, we feed signal candidates to a graphical model that handles slight signal shifts and clustered signals. The graphical model connects signal candidates across sequencing cycles by combining probability predictions and spatial distances (**Fig 1a, and Online Methods**), and resolves the sequences providing final barcodes.

**Figure 1.**
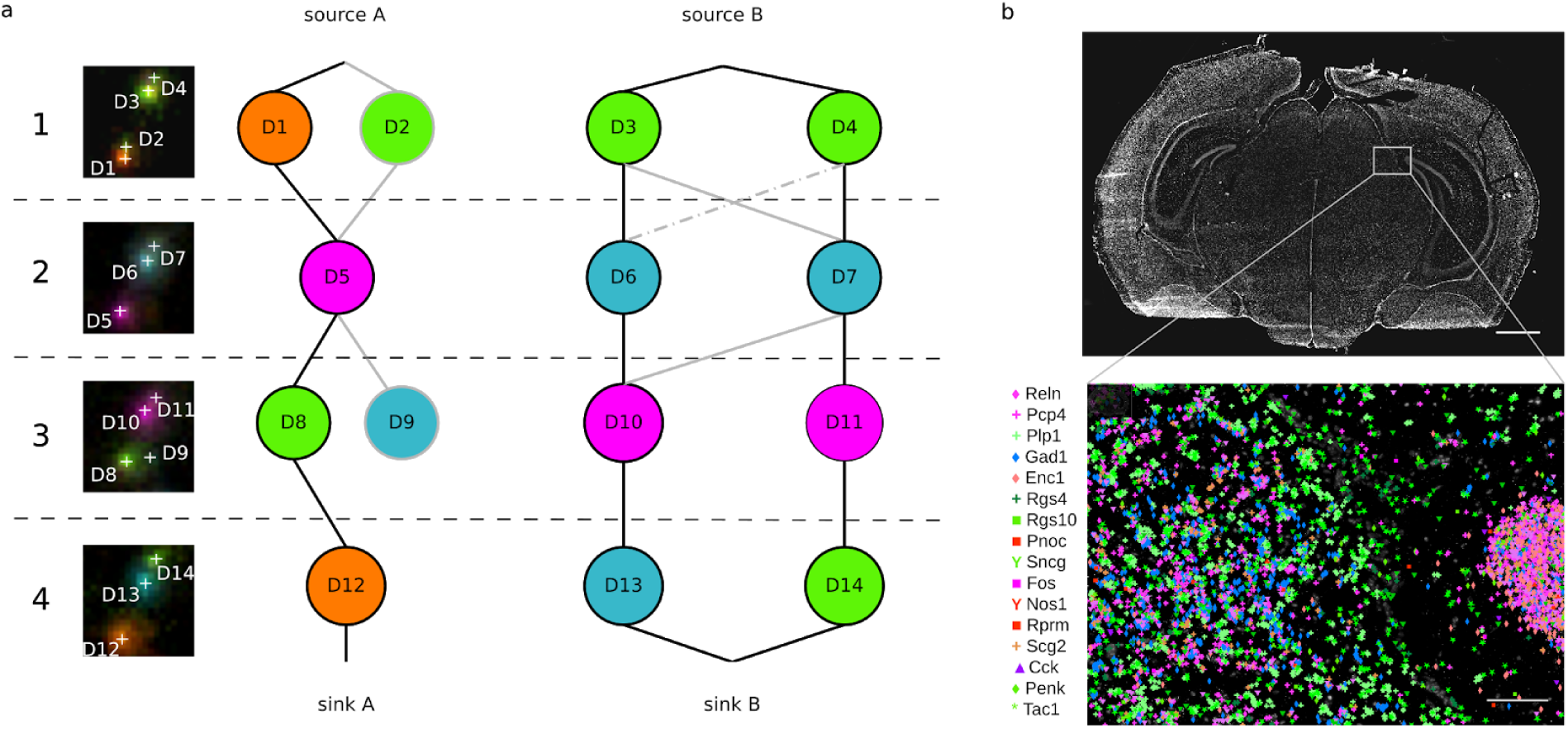
Graphical model representation for in situ sequence decoding and spatial mapping. **a)** Cut-out composite images from four sequencing cycles with four fluorescent channels (magenta, cyan, orange and green representing the letters A, C, T and G) are shown in the left panel. Each detected signal is marked with a white cross, labeled (D1-D14), and represented as a node in the graphical model (with the same color and label). The graphical representation of the fluorescent signals results in two independent connected components, represented by two graphs with label A and B. Edges between nodes represent distance connections between signal detections, where bold lines represents direct connections (distance < *d*_*th*_), and dash-dotted lines represent forced connections (distance < *d*_*max*_). Solving the graph will decode the sequences highlighted in the black paths: {D1,D5,D8,D12}, {D3,D6,D10,D13}, {D4,D7,D11,D14}, corresponding to TAGT, GCAC and GCAG. **b)** Overview of a full coronal section of a mouse brain with a zoomed-in region showing detected barcodes and corresponding genes. Scale bar top: 1000 μm, bottom: ∼100 μm.

The graphical model can either search for signals using a priori information on barcode designs, or search blindly, providing feedback on potential molecular variability. Finally, a novel sequence quality measurement is defined, making it possible to set a threshold for the desired precision-recall ratio. An example of achieved signal detection density is shown in **Fig. 1b** and a full sample is available for interactive viewing at https://tissuumaps.research.it.uu.se/demo/isseq.html, as described in **Supplementary Video**.

As in many microscopy applications, ground-truth data for methods evaluation is difficult to obtain unless one has access to alternative detection methods. We evaluated our signal decoding approach on mouse brain sections by comparing decoding results obtained at two different resolutions – thus pushing the signal size and signal-to-noise ratio of detection at the lower resolution samples. We also applied state-of-the-art in situ sequencing decoding pipeline^7^ to the same sample, and compared precision and recall of decoded sequences with respect to the proposed graph-based approach using decoded high resolution data as ground truth. We further verified the output of the decoded gene expression patterns from mouse brain sections with the corresponding patterns from Allen Mouse Brain In Situ Hybridization Atlas^1^ using Kullback-Leibler divergence (**Supplementary Text, Supplementary Fig.1**). Many expression patterns are highly correlated, while others, although visually consistent, match less accurately due to differences in tissue shapes and sizes, or strong misalignments of compared brain structures (Supplementary Fig. 2, 3).

With the rich output achieved by in situ sequencing, the next step was to gain an understanding of the spatial organization of tissue samples based on gene expression patterns, and correlate these patterns with local tissue morphology. We analyzed imaging data from in situ sequencing experiments for 97 and 82 targeted genes (listed in **Supplementary Table 2**) and decoded a total of 1955408 and 1202760 targeted barcodes per mouse brain slice (**Fig. 2a**, dimensions are respectively 10.85 mm × 7.26 mm × 9.6 μm and 10.25 mm × 7.26 mm × 8.4 μm for left and right brain). We first visualized gene expression variations across two coronal mouse brain sections mapping the gene expression profiles extracted from overlapping patches in a 3D space by UMAP manifold learning technique for dimensionality reduction^13^ (Supplementary Text). This allowed us to visualize the gene expression variation between brain regions and between samples, and the gradual shading of one gene expression profile into another between adjacent brain regions (**Fig. 2b**, Supplementary Fig. 4). Next, we explored three different techniques to extract significant expression patterns and define regions that identify brain compartments based on the expression profiles of each overlapping patch (Supplementary Text). We applied: (1) Non-Negative Matrix Factorization^14^, a linear dimensionality reduction technique, that provides a series of locally correlated gene expression maps representing distinct biological processes (Supplementary Fig. 5). (2) SpatialDE^15^ identified spatially variable genes and clustered their spatial profiles based on a Gaussian-Process-based prior. Genes with similar spatial variation are thus grouped together into spatially significant patterns (Supplementary Fig. 6). (3) UMAP dimensionality reduction followed by density based clustering of gene expression profiles of each patch. Each patch was thereafter color-coded according to the clustering and projected back to its spatial location (**Fig. 2c** and **d**, Supplementary Fig. 7). We further investigated structural variation in the gene expression between regions within and between samples. First, we defined regional divisions of brain tissue by clustering the gene expression of a set of well-known marker genes. Next, we made a differential expression analysis for all the other genes targeted by in situ sequencing (Supplementary Text, Supplementary Fig. 8).

**Figure 2.**
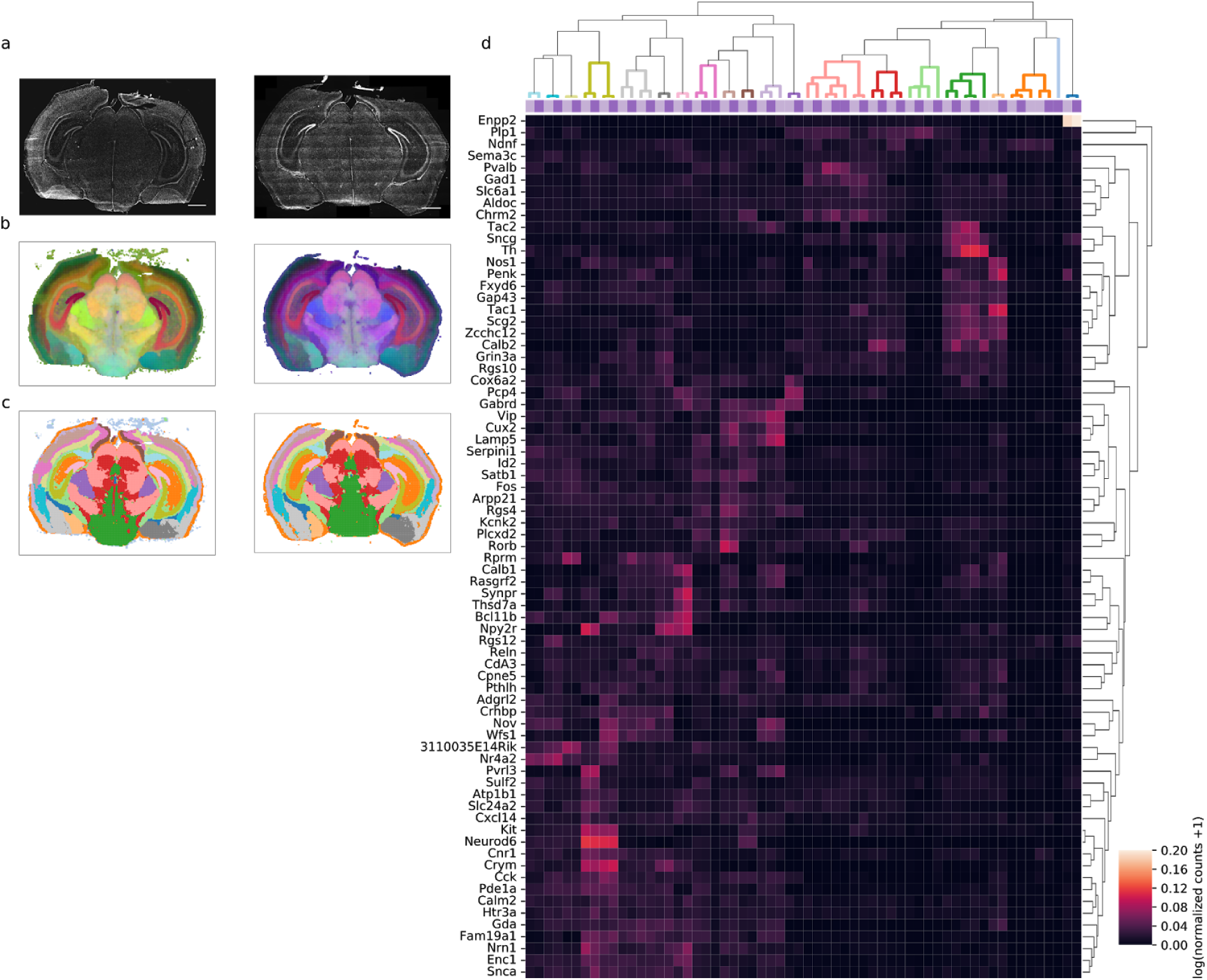
Spatial gene expression of mouse brain coronal sections. **(a)** Nuclei channels of two coronal mouse brain sections. Scale bars: 1000 μm. **(b)** Spatial gene expression variation. Patches with similar color have highly correlated gene expression profiles while color gradients indicate gradients in gene expression. **(c)** Color coded clusters of gene expression patterns on brain sections from two different mice. **(d)** Hierarchical clustering of gene expression profiles within the 20 regions from (c), where each region profile is defined by summing the expression of all patches belonging to the same region and brain, and normalizing by region size. Color codes as in (c). Profiles from corresponding regions of the two brain sections from a) cluster together (dark purple = left brain, light purple = right brain).

We further explored how local cell morphology correlates with regions defined based on gene expression patterns. Using CellProfiler^16^, we extracted 16 cell features from the nuclei staining of the sequencing data, describing cell neighborhood relationships, shape, intensity and texture (Supplementary Text). We then clustered each region based on the morphological features extracted from the same-sized patches as used for gene expression analysis (Supplementary Fig. 9). While some regions show distinct morphological profiles (i.e. cluster H and D of Supplementary Fig. 9, corresponding to CA1 and CA3 pyramidal layers, and clusters A-J-K-P, corresponding to iso-cortex), others can be better identified based on their gene expression (i.e. cluster E-B-G of Supplementary Fig. 9), since variations in cell morphology are very small.

## CONCLUSION

To conclude, we proposed a computational framework for efficient and accurate decoding of in situ sequencing 2D and 3D data in dense and noisy whole tissue samples. The proposed pipeline achieves higher recall of decoded barcodes without losing precision with respect to the state-of-the-art image analysis pipeline (Supplementary Fig. 10). The achieved decoding resolution allowed us to study the spatial organization of mouse brain coronal sections with high accuracy and reproducibility. Analysis of decoded gene expression profiles revealed biological processes driven by genes with correlated spatial expression patterns, and identified the spatial organization of different brain regions. We further showed that identification of brain compartments based on gene expression features has more discriminative power than morphological feature analysis, and thus can better guide the identification of spatial regions for further investigations in relation to development or disease.

## ONLINE METHODS

### In situ sequencing data generation

The design and experimental details of the in situ sequencing assay used to produce the data for this paper is described in detail in Qian et al.^17^. At each sequencing round, the tissue sample is stained and imaged in six fluorescent channels: a nuclei channel, a general stain channel (used as reference channel for each barcode probe), and four color channels one for each letter of the genetic code (A, C, G, T), following the protocol described by Ke et al^7^.

### Image Decoding Pipeline

The result of the experimental procedure is image data with three spatial dimensions (x, y, z), one color dimension encoding the different fluorescent channels, and a temporal dimension for the different imaging/sequencing cycles. Below we describe the analysis pipeline. The output is the spatial coordinates of decoded barcodes along with scores to assess the quality of the decoding.

### Image Registration and Tiling

Images are first aligned to compensate (i) for chromatic aberration of fluorescent channels and (ii) for misalignments among successive imaging cycles caused by repetitive washing and staining procedures. A registration step performs a coarse alignment on downsampled images of the whole tissue slide with multiresolution image registration^18^ using either rigid or affine transformation in order to preserve tissue structures. Each color channel is aligned to the general stain image of the same round to compensate for chromatic aberration. Successively, images from different rounds are aligned to a common reference sequencing cycle in order to create a common coordinate space. Therefore, for each sequencing round a transformation matrix is estimated from the registration of the general stain (or maximum intensity projection of the general stain and the nuclei channels) and a respective image of the sequencing round chosen as reference. The estimated transformation matrix is then applied to all the channels of the same round. After registration of whole slide images, each image is tiled in smaller non-overlapping patches of 1028×1028 pixel size in order to split the dataset in smaller multidimensional tensors used for parallelization of later operations and optimize memory resources.

A second alignment is then performed for each individual tensor to locally align channels and rounds repeating the procedure of the first alignment.

### Image Normalization

Images are then normalized scaling the intensity values between the background level and the signal level estimated from *n* random 128×128px patches from the whole slide image of the respective channel and round as follows:

- background level: mean value of the patch modes,
- signal level: 99th percentile of 98th patch percentiles.

### Signal Candidate Detection

Signal candidates are extracted with an h-maxima transform^19^ from the normalized images after a top-hat filtering used for enhancing bright spots and attenuate background. Therefore, all local maxima with a dynamic higher than a given threshold *h* are considered as signal candidates.

### Signals Merging

Due to the broad emission spectra or imperfect washing procedures, fluorescent signals can bleed-through to adjacent channels or rounds, and cause multiple false detections of the same signal. According to the nature of the target signal, each barcode can be fluorescent at every sequencing cycle in a single color channel other than the general stain. Therefore, signal candidates of a given sequencing round are grouped across color channels such that overlapping detection or detections that are adjacent in a 4 connectivity pixel grid are merged together keeping the signal candidate in the channel with highest intensity.

### Signal Candidate Predictions

The probability prediction of each signal candidate of being signal and noise are assessed by a convolution neural network (Supplementary Fig. 11), implemented and trained in-house on a subset of annotated candidates (nominated from the previous steps) from multiple in situ sequencing experiments. Using 5×5 px windows centered in each signal candidate (selected randomly across all color channels and sequencing cycles) as training data, the CNN learns the underlying discriminative features to predict the similarity between a signal candidate and a true signal.

### Graph-based Signal Decoding

Signal candidates and probability predictions are then combined in a graphical model to resolve and decode the final sequences across fluorescent channels and sequencing rounds. Signal candidate detections are represented in the graph as *D* nodes (colored nodes in Fig. 1a). Each *D* node consists of a pair of nodes connected by an edge with weight *w*_*i*_ equal to:

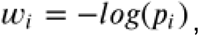

where *p*_*i*_ is the probability prediction of the signal candidate detection *i*. Relationships among signal candidate detections belonging to different sequencing cycles are encoded as edges connecting *D* nodes (Fig. 1a). Each edge connecting a pair of *D* nodes has a weight proportional to the euclidean distance between the signal candidate detections represented, specifically:

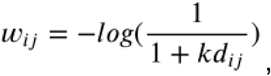

where *d*_*ij*_ is the euclidean distance between detection *i* and *j*, and *k* is a weighting parameter used to modulate the contribution of *d*_*ij*_.

In order to build the graph, signal candidate detections are searched for connected components between sequencing rounds within a maximum connection distance *d*_*th*_. Each connected component can be represented as a graph with *D* nodes encoding candidate detections and edges representing connections. Each of the connected components that are found is then refined by adding edges between not connected *D* nodes belonging to consecutive sequencing rounds that are closer than a maximum distance *d*_*max*_. Nodes of the first and last sequencing rounds are then connected respectively to a source and a sink node. Finally, connections are removed between detections belonging to not consecutive sequencing rounds and the graph is solved by maximum flow of minimum costs between the sink and the source^20^.

### Quality of Decoded Barcodes

A quality metric *Q*_*s*_ is assessed for each decoded sequence *s*, encoded by the set of detections *D*_*sh*_ :h ∈ [1,*n*] where *n* is the number of sequencing cycles. The quality score per sequence is proportional to the probability predictions, intensities and distances of the signal candidate detections that form the sequence, and is defined as:

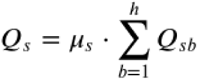

where *Q*_*sb*_ is the quality score of each decoded base that form the sequence and *μ*_*s*_ is a function proportional to the maximum distance between the detections that form the sequence. Specifically *Q*_*sb*_ is defined as:

- if multiple detections {*D*_*sb1*_,…,*D*_*sbk*_}, with a probability prediction higher than 0.5, other than *D*_*sb*_ where detected and merged in a given round :

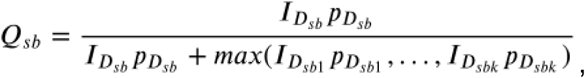
- otherwise,

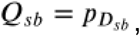

where *I* and *p* are respectively intensity value and probability prediction of a given candidate. In order to penalize sequences whose detections are far apart from each other, *μ*_*s*_ is defined as:

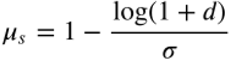

and clipped between 0 and 1 values, where *d* is the maximum distance between signal candidate detections composing a sequence and *σ* is a parameter weighting the penalty. Parameter *σ* can be empirically set to the value that will maximize the Area Under the Receiver Operating Characteristics (Supplementary Fig. 10) from the true positive rate and false positive rate evaluation based on sequences of targeted barcodes.

### 2D vs 3D Analysis

The proposed image analysis pipeline allows to resolve and decode barcodes in 2D and in 3D. Where, 2D analysis of maximum projected images of 3D focal planes are usually preferred for faster processing time. Whereas detect and resolve fluorescent signals in 3D can better disentangle overlapping signals and dense clustered regions, thus achieving better recall. A comparison between 2D and 3D analysis results is shown in Supplementary Fig. 10.

### Code Availability

All software was developed in Python 3 using open source libraries, and data processing of pipeline workflows were carried out using Anduril2 analysis framework^21^. The processing pipeline and the software version used to generate the analysis results presented in this paper are available at https://github.com/wahlby-lab/graph-iss.

## ACKNOWLEDGMENTS

We thank Xiaoyan Qian for kindly providing in situ sequencing images and Fred Hamprecht for valuable input on signal decoding. This research was funded by the European Research Council via ERC Consolidator grant 682810 to C. Wählby.

## AUTHOR CONTRIBUTIONS

GP provided the main contribution to design, implementation and evaluation of image decoding pipeline, spatial gene expression analyses, figure preparation and manuscript text. GP, CW and LS contributed to morphological feature analysis. MH contributed to in situ sequencing experiments, design of spatial gene expression analysis, manuscript text and figures. GM contributed to the design and implementation of image decoding pipeline. LS developed the online viewer application allowing visual examination of results. AK contributed to evaluation of the image decoding pipeline, produced decoding results for the state-of-art pipeline, and contributed to figure preparation. MN contributed to the design of spatial gene expression analysis. CW and MN contributed to the conception of the study. CW contributed to manuscript text and figures, and the design of image decoding pipeline evaluation and spatial gene expression analysis.

