## Supplementary Text for "Identification of spatial compartments in tissue from *in situ* sequencing data"

### Datasets

In situ sequencing images (acquired with Zeiss ZEN software) were from the recent study by Qian et al[1]. In brief, 10 µm fresh frozen brain tissues from a CD1 male mouse (postnatal day 25) were probed with 95 nucleotide (nt) long padlock probes. Padlock probes contained a 4 nt sequencing by ligation barcode and a 1 nt sequencing by hybridization barcode. A 99-gene panel and a 84-gene overlapping panel (**Supplementary Table 2**) were probed in two different experiments and imaged on an epifluorescence microscope AxioImager.Z2 (Zeiss) with 20x/0.8 objective. Respectively, two genes (*Npy* and *Sst*) of the 99 and 84-gene panels were detected separately in 6th hybridization round. One set of images of each experiment (channels: DAPI, FITC, Cy3, TexasRed, Cy5 and Cy7) was decoded without the 6th sequencing round and analyzed as described below.

### Spatial Gene Expression Analysis

We investigate the spatial gene expression variation of 82 genes targeted in 5 sequencing rounds in both mouse coronal sections with three different clustering techniques. Reads are first decoded with the described image analysis pipeline and filtered to remove low quality reads applying a quality threshold of 2, and excluding low expressed genes that have a total count lower than 500 reads. Successively, a gene expression matrix is constructed for each sample where rows represent overlapping patches of the tissue sample and columns represent targeted genes. Gene expression matrices are generated from the two sections using square tiles of size 128 px and overlap 128 px for a total patch size of 384x384 px. Each entry in the matrix represents the expression level of a particular gene in a given patch (i.e. total count of reads decoded inside the patch). An additional filtering step excludes patches with less than 10 reads from further evaluations. Thus, a total of 73 genes common to both samples passed the filtering steps, resulting in a total of 11 genes excluded from the analysis (i.e. *Chodl*, *Cort*, *Crh*, *Hapln1*, *Lhx6*, *Npy*, *Pax6*, *Qrfpr*, *Rab3c*, *Slc17a8*, *Sst*). The expression table of each sample is then normalized (as in Svensson et al.[2]) stabilizing variance using Anscombe's transform and successively regressing out the logarithmic total counts of reads per patch in order to exclude unwanted sources of variation such that, number and size of cells in each patch. We finally scale the expression matrices feature-wise to zero mean and unit variance.

*Spatial Gene Expression Gradient*

First we inspect gene expression spatial variation across the coronal sections, visualizing gradient of gene expression profiles between brain regions (Fig. 2b, and Supplementary Fig. 4). Each of the filtered and normalized expression matrices are mapped into a common 3-dimensional embedding using UMAP[3]. The axis of the 3-dimensional embedding are then normalized to unit vectors and coordinates of each projected patch in this new reference space are used to map each patch in the RGB color space.

*Spatial Gene Expression Clusters*

We then investigate if the gene expression from the two sections clusters in common brain regions. The gene expression matrices generated in the previous analysis are separately mapped into a 10-dimensional space using UMAP and clustered using

Bayesian Gaussian Mixture Model[4]. Initializing the number of mixture components to 30, visually coherent clusters of anatomical structures appear in both brains characterized by highly correlated gene expression signatures (Fig 2c-d, Supplementary Fig. 7).

In order to perform differential expression analysis between regions without bias caused by the clustering itself, we define regions by clustering the gene expression of 18 marker genes. Marker genes were chosen hierarchically according to strongly expressed genes for interneurons, pyramidal cells and non-neuronal cells[1].

We first apply a dimensionality reduction of the marker gene expression profiles of the two brains in a common lower dimensional space. We then cluster in 20 regions each brain individually with variational bayesian estimation of gaussian mixtures, using the solution of the first fit as initialization for the second brain (Supplementary Fig. 8a-b). We now perform differential expression analysis with Seurat[5] for the rest of the 73 genes between and within the defined regions in the two brains that have a sample pairwise correlation higher than 0.9 (Supplementary Fig. 8c-d).

*Non Negative Matrix Factorization*

Non negative matrix factorization is well suited for its non-negativity feature to biological contexts where biological signals can be naturally present or absent.

We therefore translate normalized expression values into a positive space for applying non negative matrix factorization analysis[6]. The expression matrix **E** is thus factorized into two nonnegative matrices **W** and **H** initializing the number of factors to 20. An approximation of the gene expression matrix E is then reconstructed as a linear combination of column feature vectors in H weighted by the contribution of each gene column in matrix W.

$$E = WH$$

Columns feature vectors of matrix H represent therefore co-expression patterns (also called metagenes) shared in the two brains, often corresponding to real biological patterns. And rows of matrix W represent the contribution of each gene in each metagene (Supplementary Fig. 5).

*SpatialDE Analysis*

We then investigate gene expression variation in tissue samples with SpatialDE[2], a framework based on Gaussian process regression that identifies genes with spatially significative gene expression patterns. We run SpatialDE analysis on normalized gene expression matrices generated from square patches of 512 px size and 64px overlap, due to a higher demand of computational resources by the tool. SpatialDE classified all targeted genes as significantly spatially varying as was expected due to the choice of targeted genes (Supplementary Fig. 6a). We then perform SpatialDE "automatic expression histology" analysis consisting in a spatial clustering of similarly spatially varying expression patterns based Gaussian Mixture Models with a spatial Guassian prior on the cluster centroids. Initializing the number of patterns to 20, we obtain 15 and 13 spatial gene expression patterns defined by sets from 1 to 24 genes (Supplementary Fig. 6b-c, and Supplementary Table 1).

### Allen Brain Atlas Comparison

We then compare in situ sequencing (ISS) decoded gene expression patterns with respect to 3d grid expression data from in-situ hybridization (ISH) Allen Mouse Brain Atlas[7]. The Allen Mouse Brain Atlas provides genome-wide in-situ hybridization data for approximately 20.000 genes processed with a data processing pipeline[8,9]. The output of the data processing is a 67x41x58 voxel grid with quantified gene expression values for each gene supporting differential and correlative spatial gene expression analyses. For each of the gene expression patterns decoded in the two brain sections we compute the Kullback-Leibler divergence with respect to grid expression patterns from the ISH atlas in order to assess spatial pattern similarities (Supplementary Fig. 1). Gene expression patterns from ISS are first scaled to match the resolution of atlas voxel grid. Secondly, a probability density function is estimated for each gene using a gaussian kernel with covariance factor of 0.05. Both ISS and ISH gene expression patterns are then normalized such that the total mass of each probability density function is 1. We then select two of the 67 coronal levels that best match the ISS sections for computing the Kullback-Leibler divergence between normalized expression patterns. The coronal level showing the closest gene-gene similarity is further selected for visualization purposes.

### Morphological Features Analysis

We then conduct morphological feature analysis to study how cell morphology relates with gene expression profiles. We first segment cell nuclei from the relative channel with CellProfiler[10] and successively we extract 16 morphological features using CellProfiler Analytics[11]. We remove border artifacts from the single tiles before normalize morphological features between bounds that visually highlight spatial profiles. And we further remove outliers based on nuclei area shape caused by segmentation errors. Morphological feature profiles are then extracted for each of the patches defined previously, and patches with less than 3 cells are excluded. We regress out the total number of cells per patch and we compare morphological profiles of spatial regions defined through gene expression clustering (Supplementary Fig. 8a, Supplementary Fig. 9)

**Quantitative Evaluation of Image Processing Pipeline**

We perform and compare analyses of approximately 1.79mm x 1.46mm mouse brain section imaged with 40x and 20x magnification objective to assess precision and recall of image processing pipelines. Specifically, we decode 40x image data in 3D and 2D with the proposed graph-based approach and 3D analysis is successively chosen as ground truth for the validation of 20x decoding analyses due to higher recall and better discriminatory power between true targeted sequences and false positives (Supplementary Fig. 10a-b). We then evaluate the results of three different decoding of 20x image data and compare them with respect to the 40x ground truth. Analysis of the already aligned 20x image data have been performed in 3D with the proposed graph based-approach, and in 2D with the proposed approach and the CellProfiler pipeline from Ke et al.[12], where parameters were manually tuned for optimal performance on the dataset. For each of the three 20x analysis results, we spatially align decoded reads to the reference ground truth applying an affine transformation estimated with SIFT[13] feature based landmark registration in Fiji[14] between maximum intensity projected general stain images of 20x and 40x first sequencing rounds. We then perform a second alignment based on locally affine point cloud registration of decoded reads. Where decoded reads are divided into patches based on their spatial location and an affine transformation is estimated and applied for each patch, using iterative-closest point algorithm with matches further than 5 pixels away excluded[1]. Aligned decoded reads are then paired with their nearest neighbor in the ground truth within a maximum euclidean distance of 3. For each paired reads the number of mismatches in their decoded sequences are then evaluated. Due to the presence in the ground truth of a consistent amount of false positive reads (caused by computational and biological noise) whose majority present one base mismatch with respect to one of targeted sequences (Supplementary Fig. 10b), we exploit the fact that targeted sequences are encoded with one base redundancy allowing to detect one base errors. Thus, in order for the evaluations to be less affected by errors in the ground truth, we consider as perfect matches (0 mismatches) reads from the evaluated analysis that present one base mismatch with the paired ground truth read and that are true targeted sequences. For each 20x analysis, we evaluate precision and recall at different quality thresholds as follows (Supplementary Fig. 10d):

$$\text{Precision} = \frac{\text{TP}}{\text{TP} + \text{FP}}, \qquad \text{Recall} = \frac{\text{TP}}{\text{TP} + \text{FN}},$$

where, for a given quality threshold:

- True Positives (TP) are reads that pass quality threshold and have 0 mismatches.
- False Positives (FP) are reads that pass quality threshold and have >0 mismatches or do not have a matching read.
- False Negatives (FN) are reads with 0 mismatches that do not pass the quality threshold, and ground truth reads that are true targeted sequences but do not have a matching read.

**On-line interactive data viewer**

An interactive viewer for sequencing data is available via the TissUUmaps project: https://tissuumaps.research.it.uu.se/demo/isseq.html. Instruction demonstrating how to use the viewer is available as **Supplementary Video**. Note that only a randomly selected fraction of the transcripts are shown at low resolution to optimize visualization interaction. Zooming in to a smaller part of the tissue will show all transcripts. Color coding and symbols and their size can be modified.

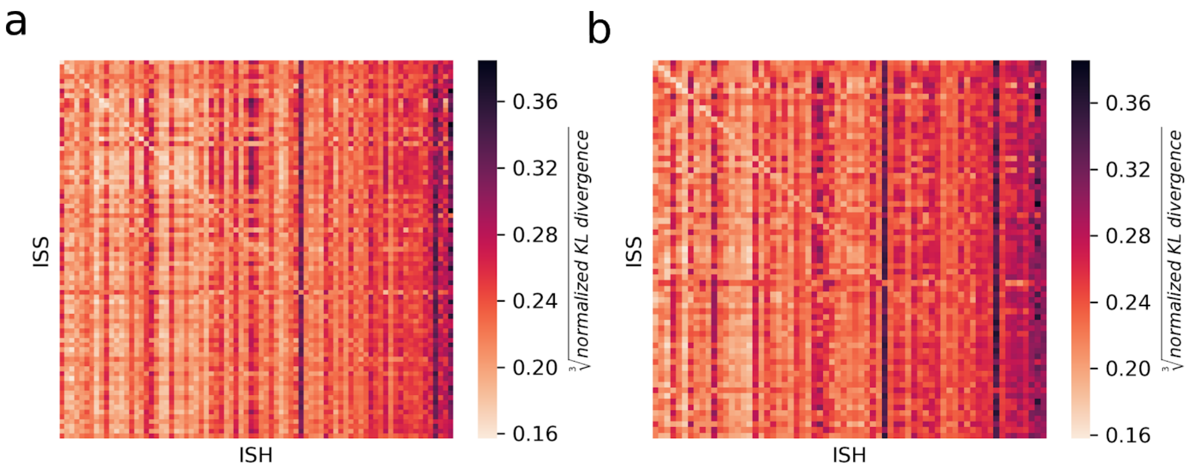


**Supplementary Fig. 1.** Allen Mouse Brain ISH-Atlas Comparison. **a,b)** Visualization of normalized KL divergence between in situ sequencing (ISS) spatial gene expression patterns and in situ hybridization patterns from Allen Mouse Brain Atlas for the left and right brain from Figure 1. Rows and columns are sorted by the difference between the KL divergence of a gene ISS pattern with its correspective ISH pattern and the minimum KL divergence with the other genes. Such that, patterns that uniquely match their pair appear in the top-left quadrant.

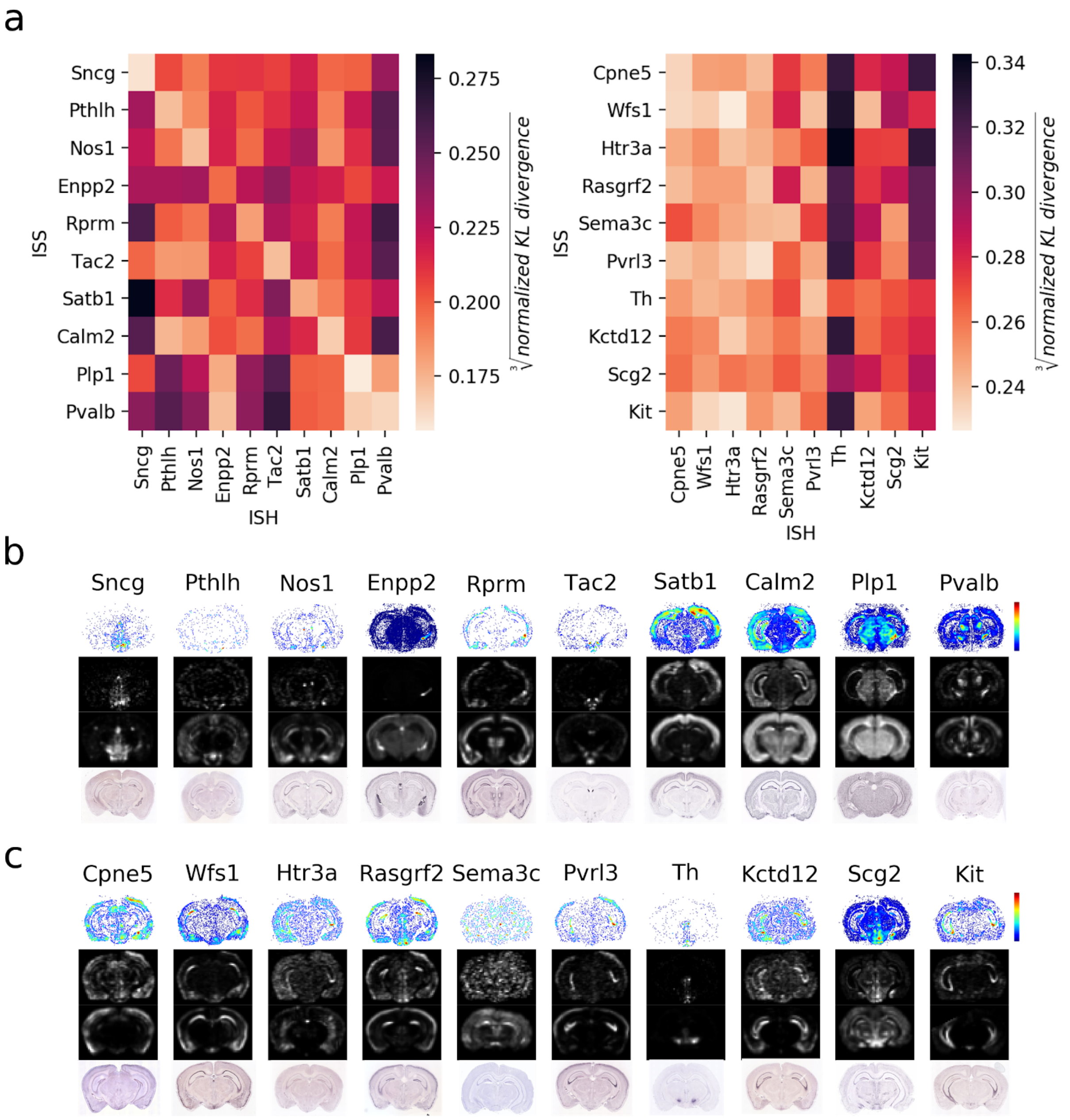


**Supplementary Fig. 2.** Detailed Allen Mouse Brain ISH-Atlas Comparison of Supplementary Fig. 1a**. a)** Detailed visualization of 10 top-left and 10 bottom-right genes of Supplementary Fig. 1a. **b)** Visualization of spatial patterns of top-left genes, from top to bottom: decoded ISS reads, generated ISS expression grid 2d histogram, Allen Brain Atlas ISH expression grid 2d histogram, Allen Brain Atlas ISH data. **c)** Visualization of spatial patterns of bottom-right genes, from top to bottom: decoded ISS reads, generated ISS expression grid 2d histogram, Allen Brain Atlas ISH expression grid 2d histogram, Allen Brain Atlas ISH data.

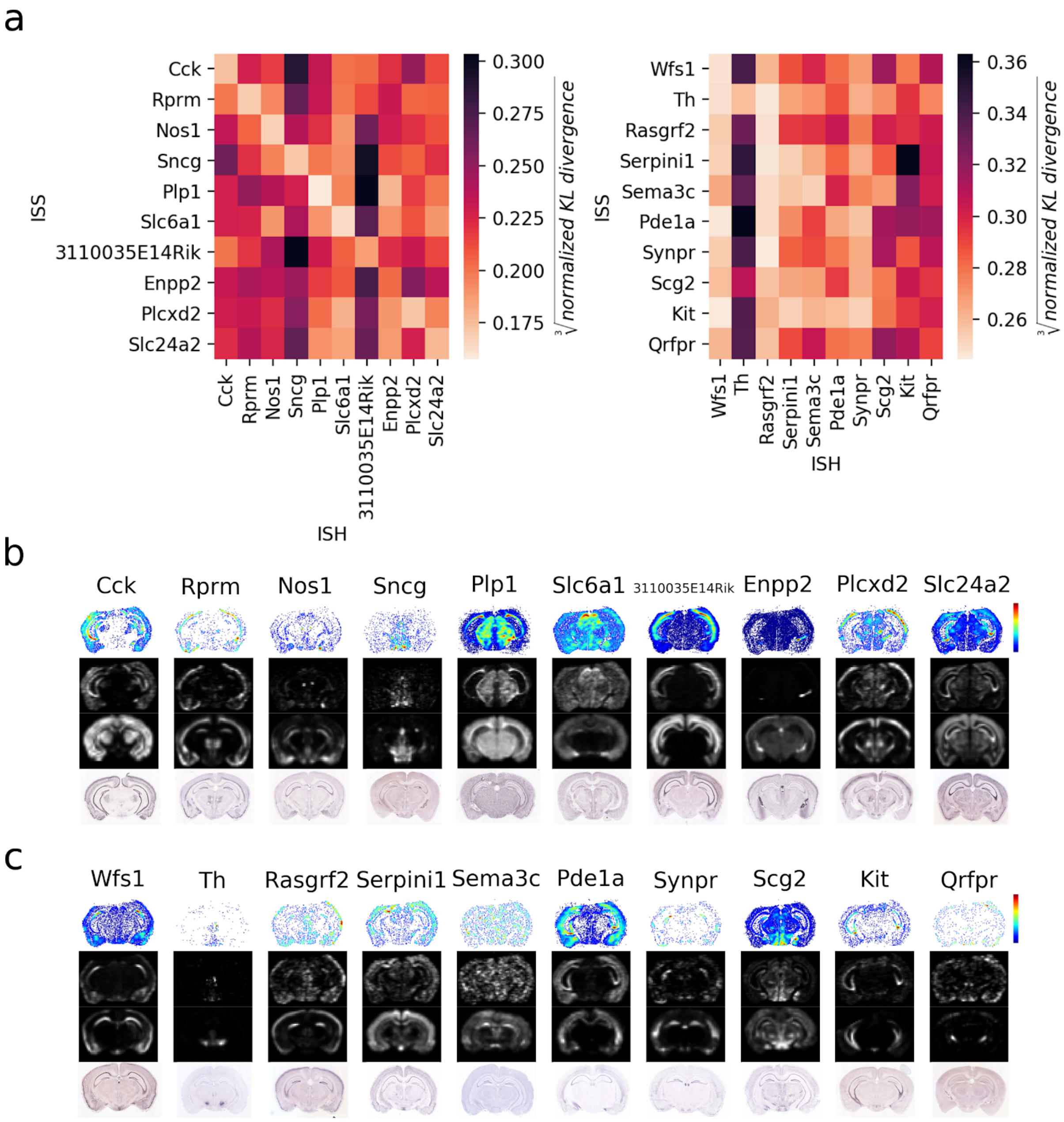


**Supplementary Fig. 3.** Detailed Allen Mouse Brain ISH-Atlas Comparison of Supplementary Fig. 1b**. a)** Detailed visualization of 10 top-left and 10 bottom-right genes of Supplementary Fig. 1b. **b)** Visualization of spatial patterns of top-left genes, from top to bottom we have: decoded ISS reads, generated ISS expression grid 2d histogram, Allen Brain Atlas ISH expression grid 2d histogram, Allen Brain Atlas ISH data. **c)** Visualization of spatial patterns of bottom-right genes, from top to bottom we have: decoded ISS reads, generated ISS expression grid 2d histogram, Allen Brain Atlas ISH expression grid 2d histogram, Allen Brain Atlas ISH data.

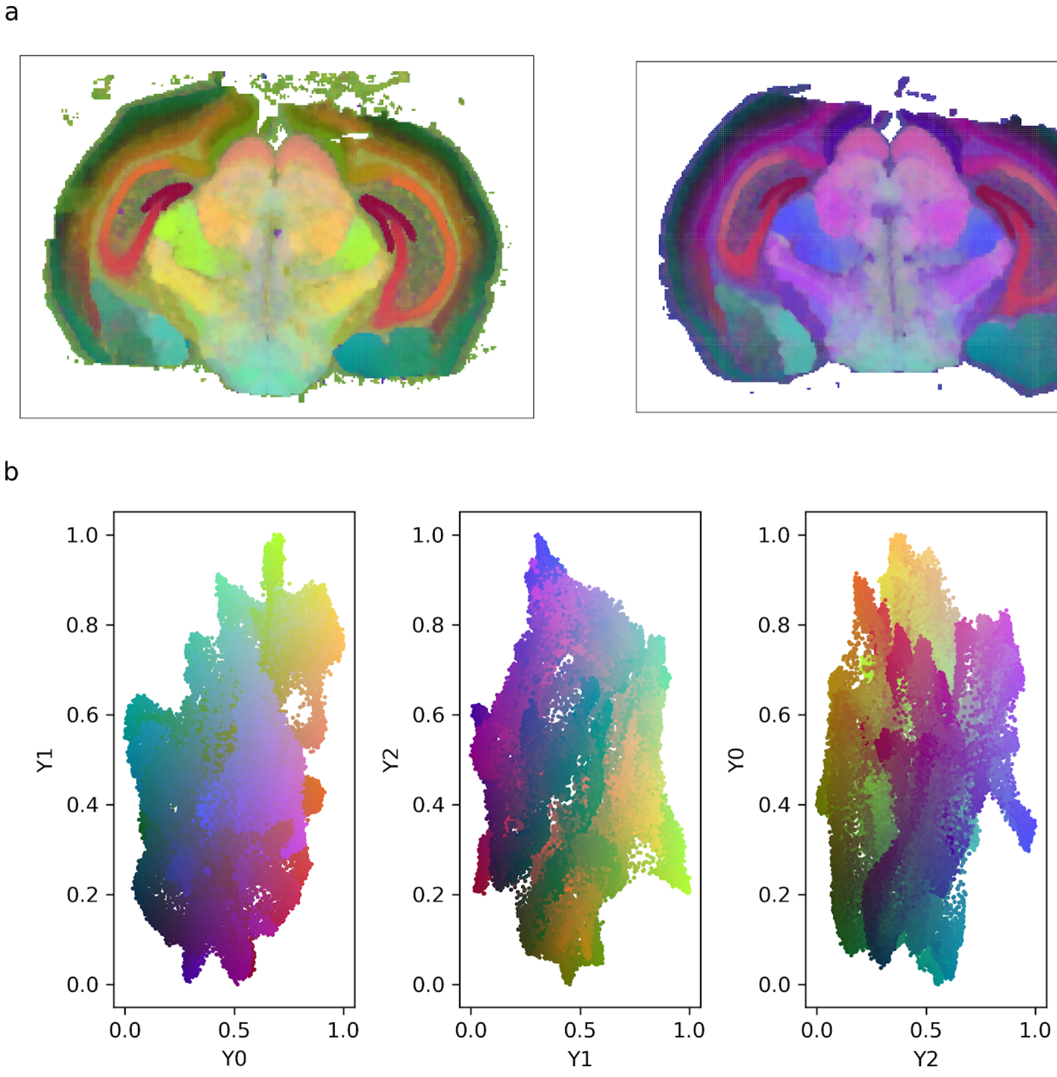


**Supplementary Fig. 4.** Spatial Gene Expression Gradient. **a)** Brain Gene expression variations. Each patch is color coded based on its gene expression profiles projected in a 3D space. Patches with similar color have highly correlated gene expression profiles. **b)** Visualization of the patch gene expression profiles in the dimensionality reduction space.

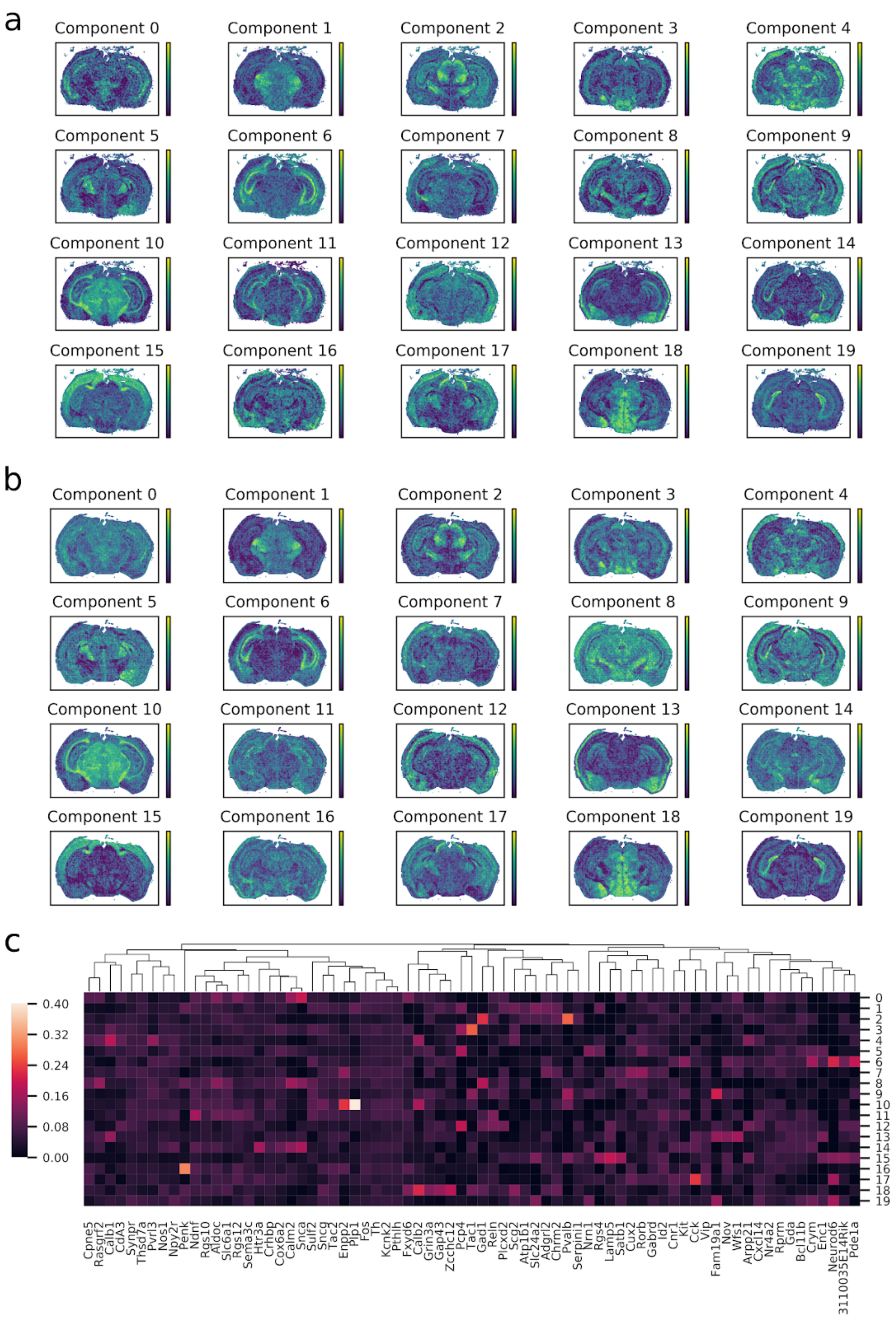


**Supplementary Fig. 5.** Non-Negative Matrix Factorization Analysis. **a,b)** Visualization of non-negative matrix factorization single components that represent co-expression patterns of the two brains. **c)** Visualization of the weight matrix representing the contribution of each gene to the different co-expression patterns.

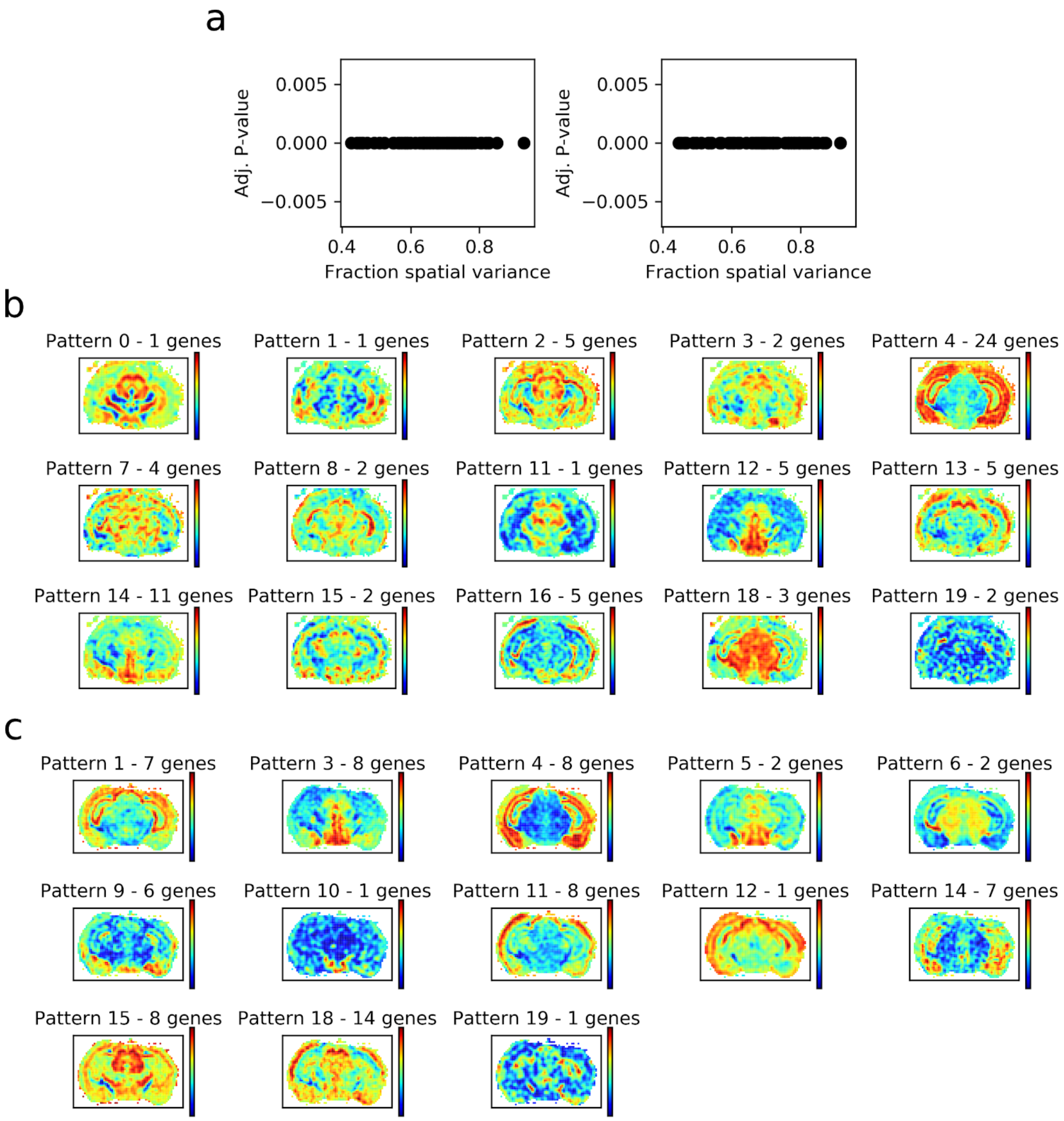


**Supplementary Fig. 6.** SpatialDE Analysis. **a)** Fraction of variance explained by spatial variation (FSV) versus significance of spatial variation for all targeted genes. **b,c)** Visualization of “Automatic Expression Histology” analysis of the two brains. Colorbars represent expression levels of each spatial pattern. The number of genes contributing to the pattern are listed in the title of each figure, and detailed in Supplementary Table 1a-b.

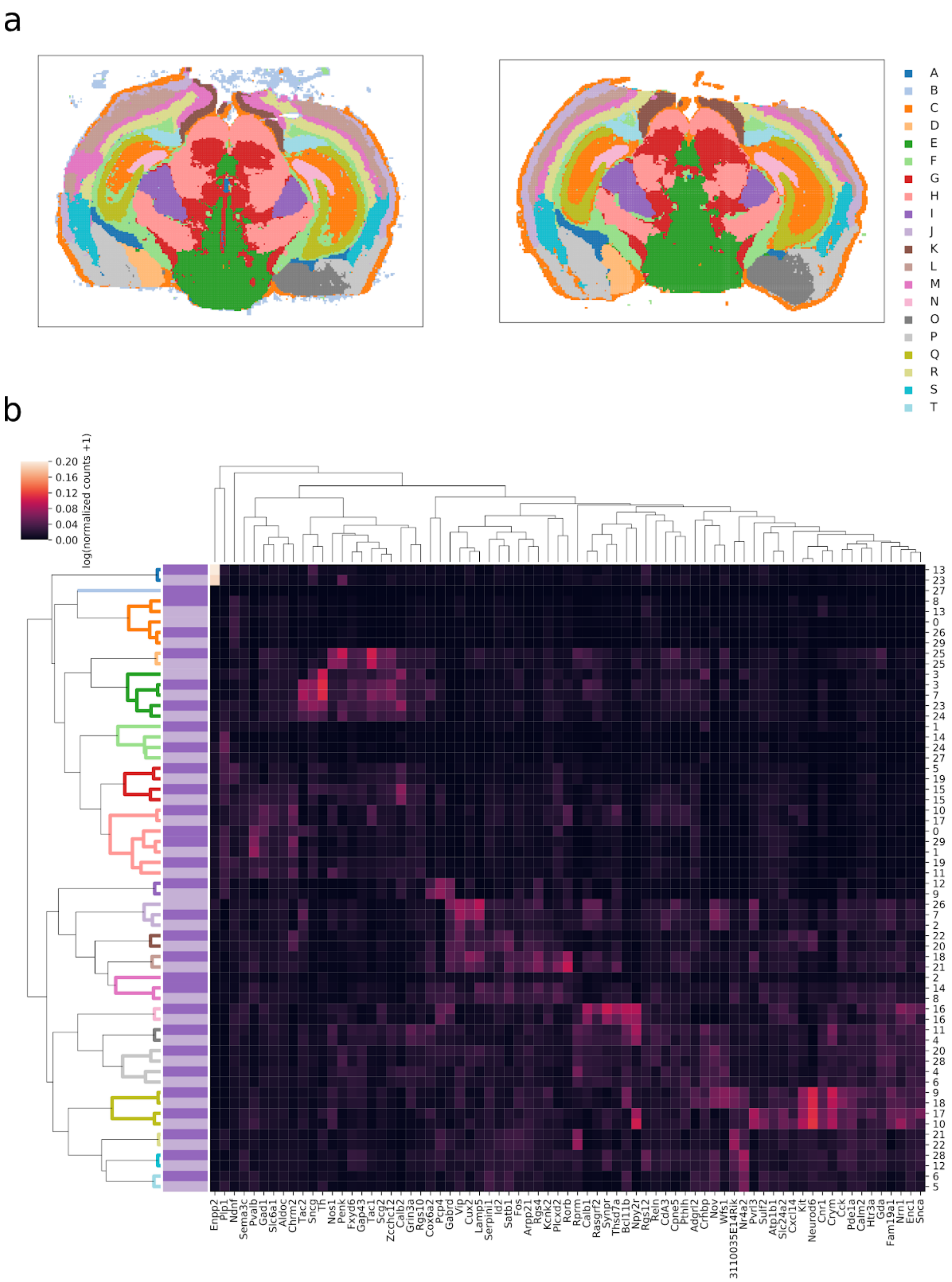


**Supplementary Fig. 7a, b.** Spatial Gene Expression Clustering. **a)** Visualization of 20 regions defined by highly correlated clusters from Supplementary Fig 7b, **b)** Hierarchical clustering of cluster gene expression profiles based on correlation. Gene expression profiles for each cluster are defined summing the expression of all patches belonging to the cluster and normalizing by cluster size. Row colors represent sample id: dark purple for left brain, light purple for right brain.

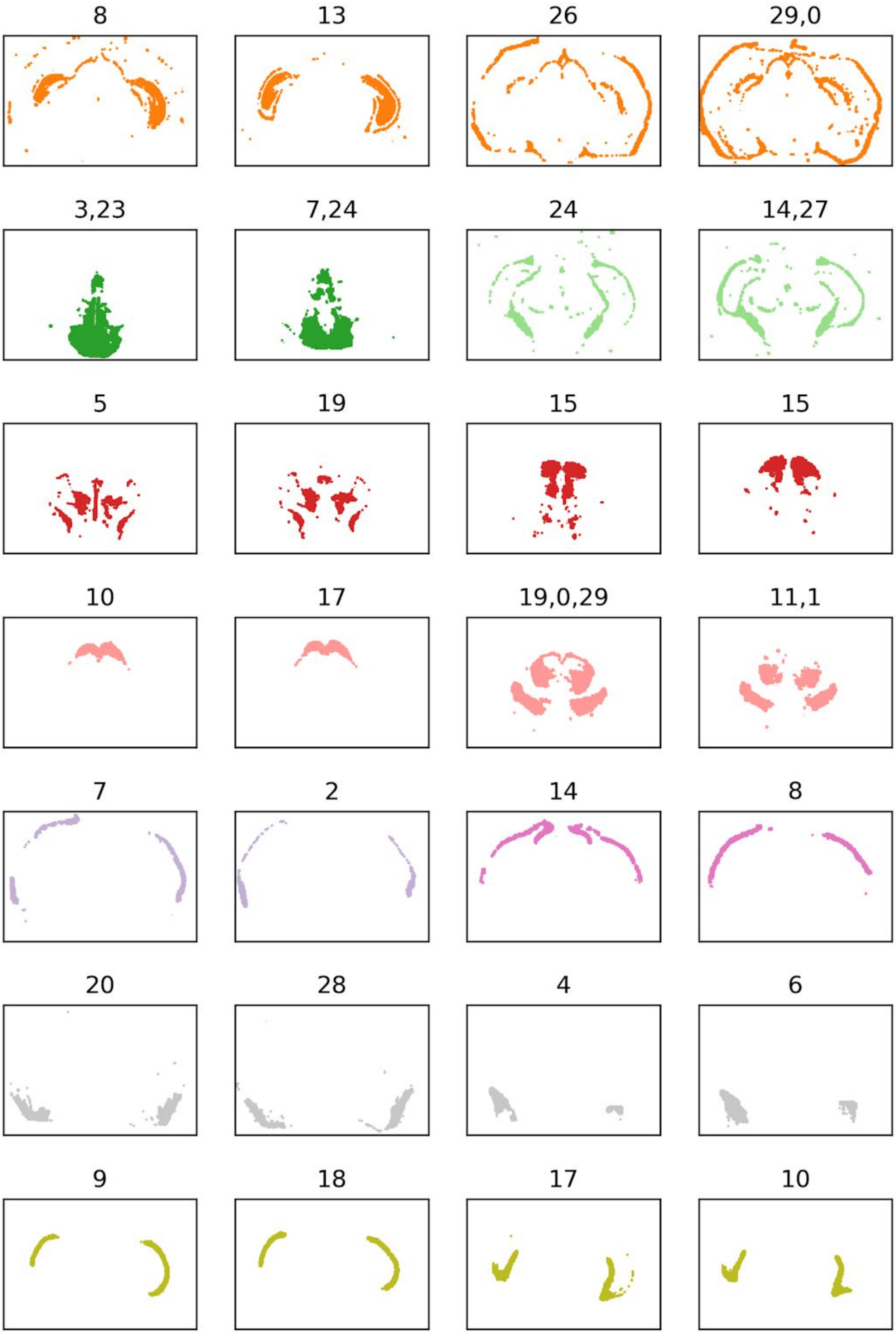


**Supplementary Fig. 7c** Visualization of pairs of gene expression clusters from the two brains. Each pair represents left and right brain of a sub-cluster of the 20 regions defined in Supplementary Fig. 7a.

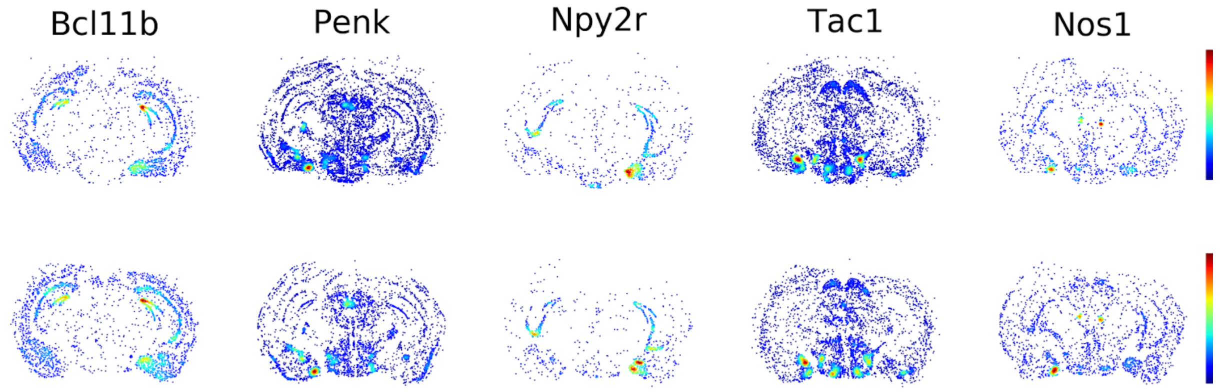


**Supplementary Fig. 7d.** Decoded reads of the most significant genes of cluster D and O from Supplementary Fig. 7a, color coded based on spatial density.

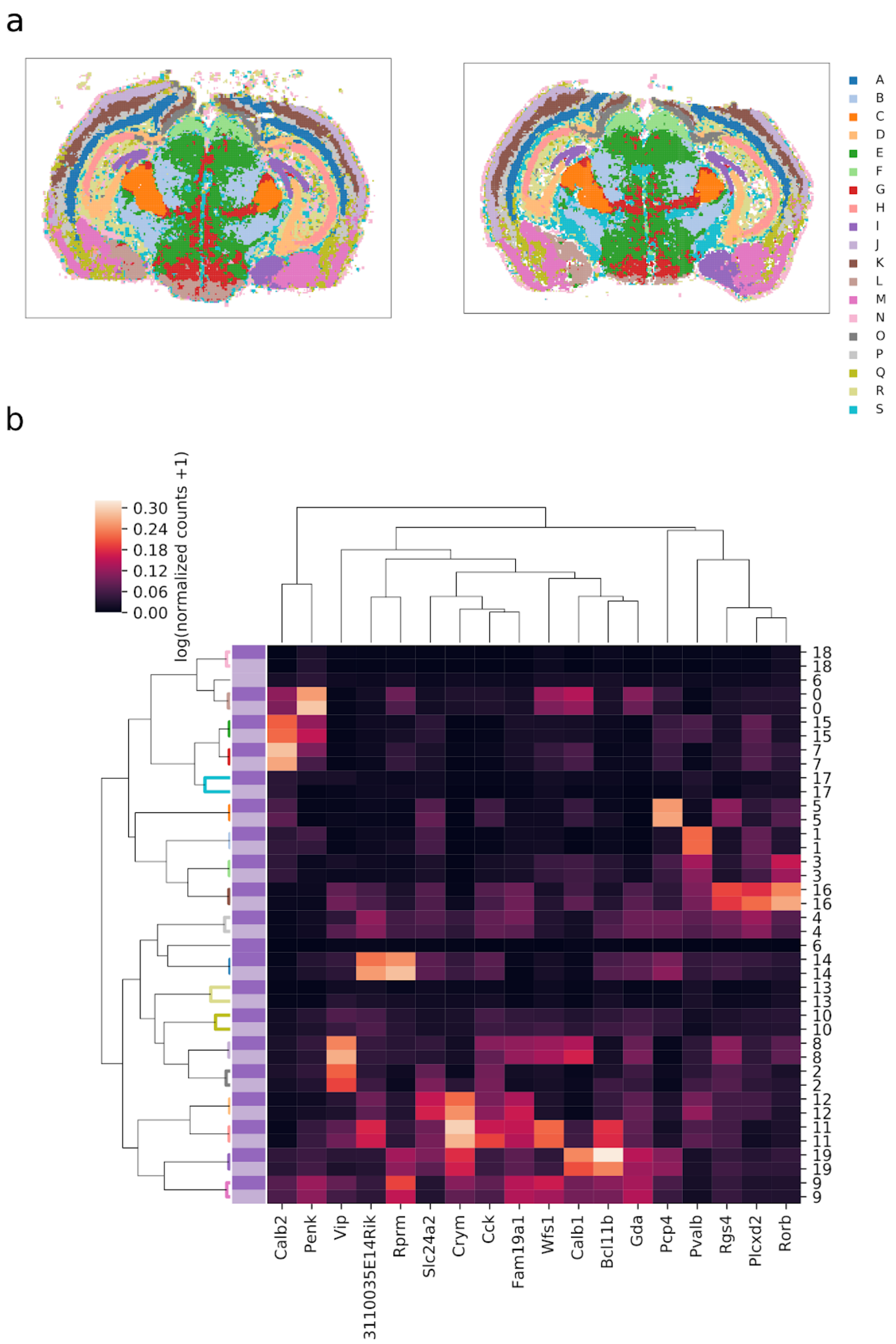


**Supplementary Fig. 8a, b.** Gene expression clustering based on 18 selected marker genes. **a)** Visualization of 19 regions defined by highly correlated clusters from Supplementary Fig. 8b, **b)** Hierarchical clustering of cluster gene expression profiles based on correlation. Gene expression profiles for each cluster are defined by summing the expression of selected marker genes from all patches belonging to the cluster and normalizing by cluster size. Row colors represent sample id: dark purple for left brain, light purple for right brain. The two rows numbered 6 represent patches with low expression counts not correlated across the two brains, and therefore not assigned a color/letter.

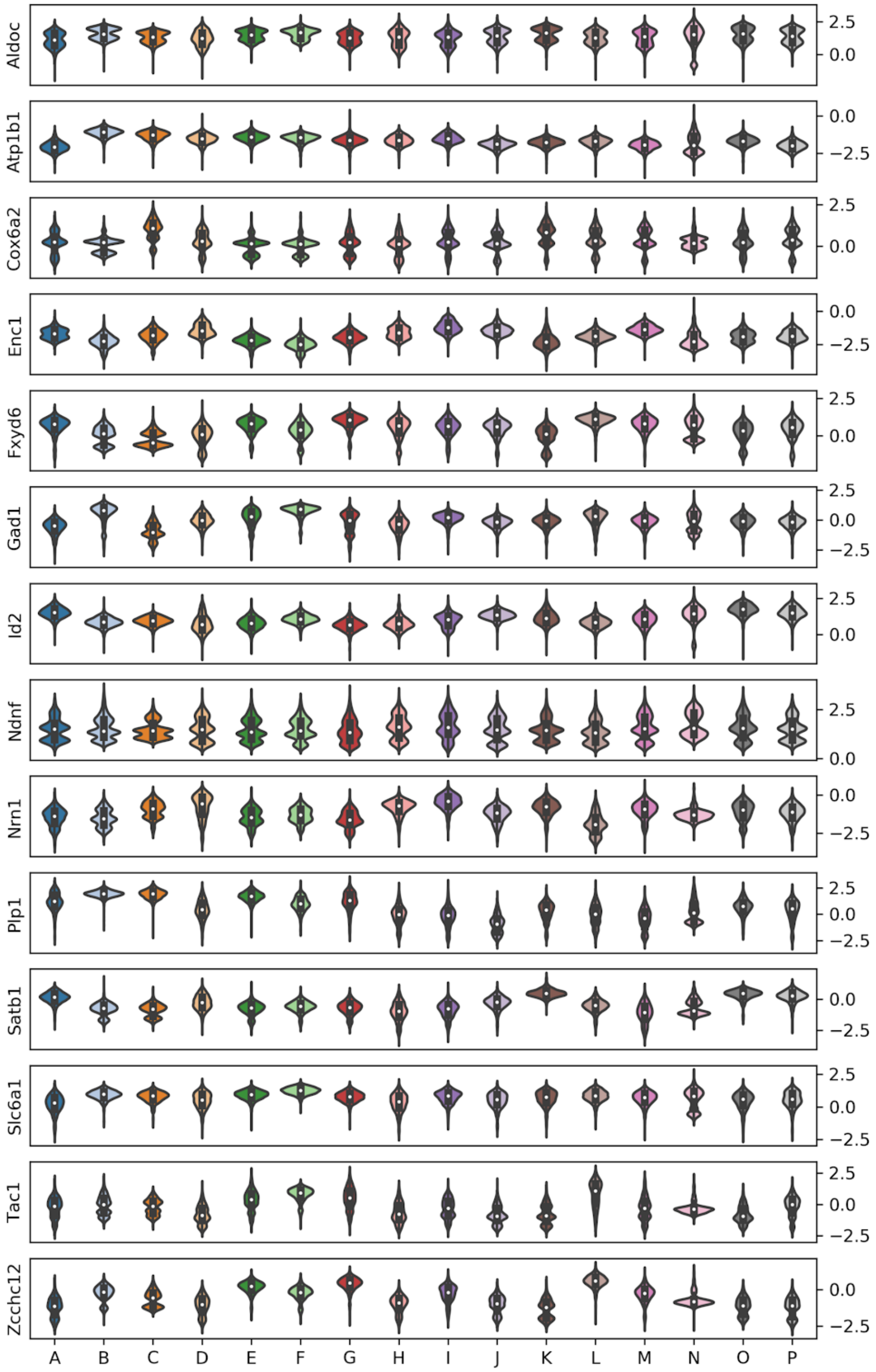


**Supplementary Fig. 8c part 1.** Differential Expression Analysis of Brain Regions. Combined normalized gene expression profiles from the two brains for each non-marker gene across each cluster defined in Supplementary Fig. 8a.

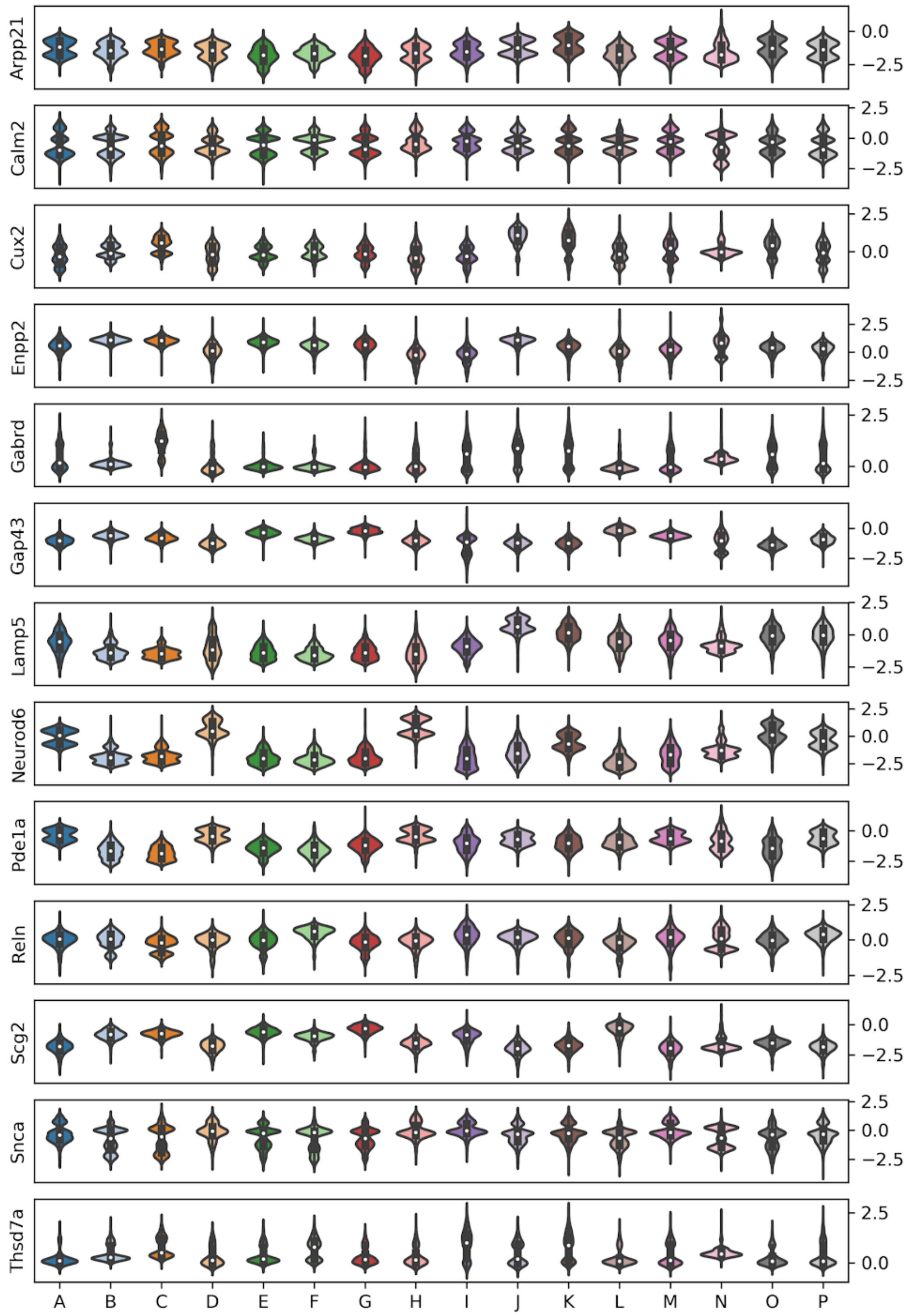


**Supplementary Fig. 8c part2.** Differential Expression Analysis of Brain Regions. Combined normalized gene expression profiles from the two brains for each non-marker gene across each cluster defined in Supplementary Fig. 8a.

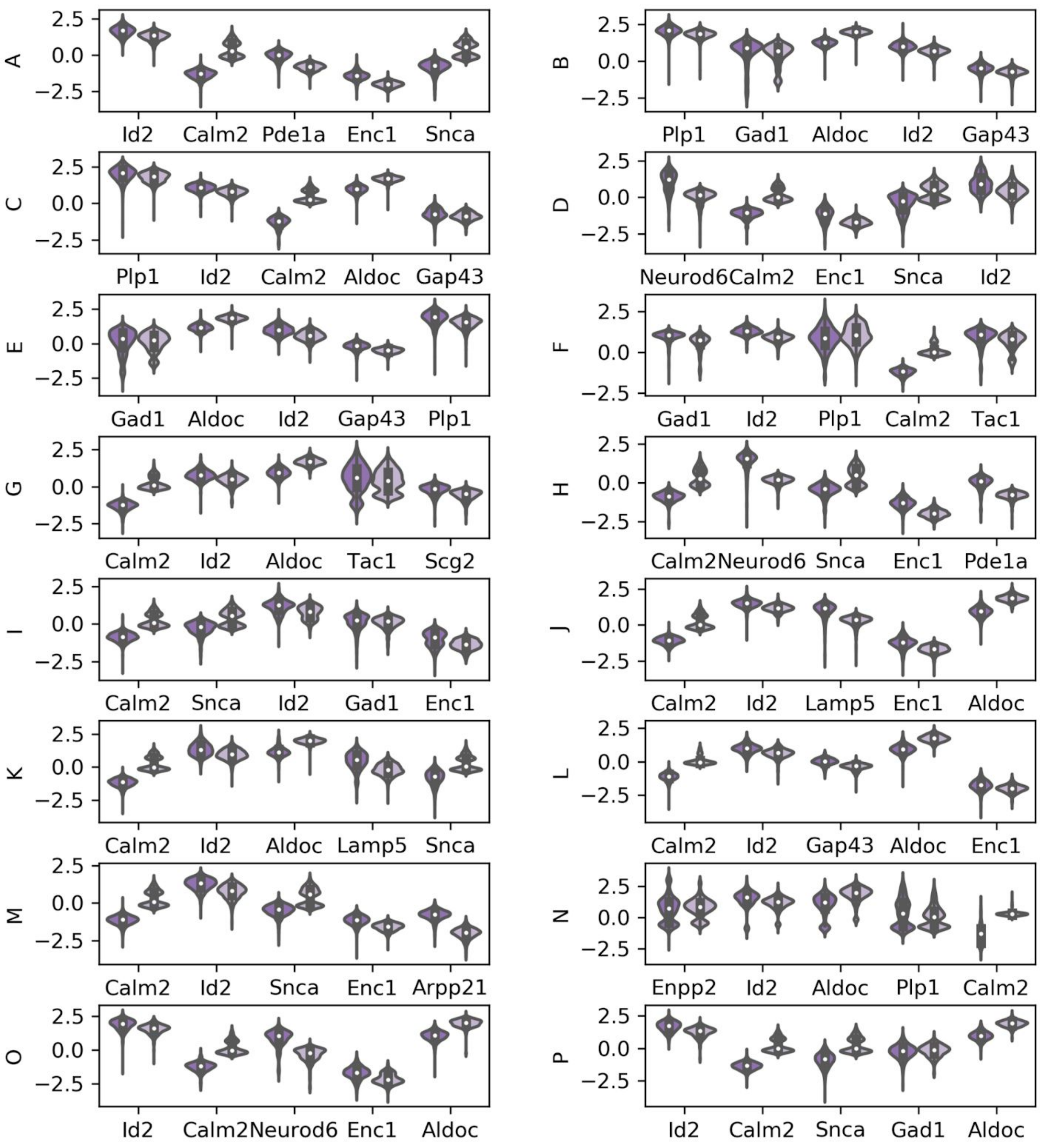


**Supplementary Fig. 8d** Cluster gene expression profiles of top five differentially expressed genes. Violin plots of normalized expression are shown for the top 5 genes based on average fold change with adjusted p-values <0.01 for each cluster defined in Supplementary Fig. 8a. Brain samples are represented with the same color code.

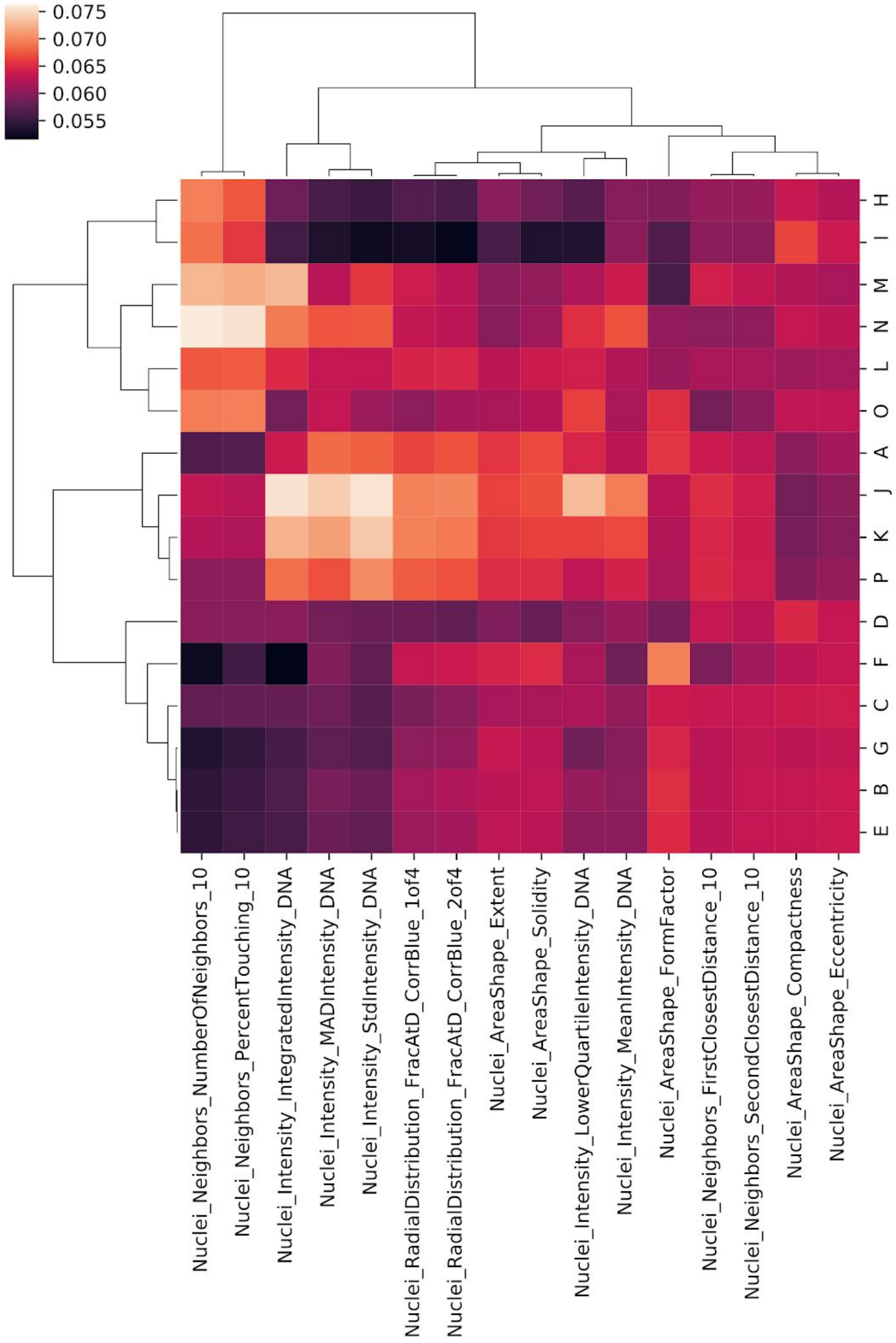


**Supplementary Fig. 9.** Morphological Feature Analysis. Hierarchical clustering based on cosine distance of morphological feature profiles of clusters defined by gene expression analysis (Supplementary Fig. 8a). Morphological profiles are defined for each cluster summing the normalized morphological features of all patches belonging to the cluster and scaling each feature by cluster size.

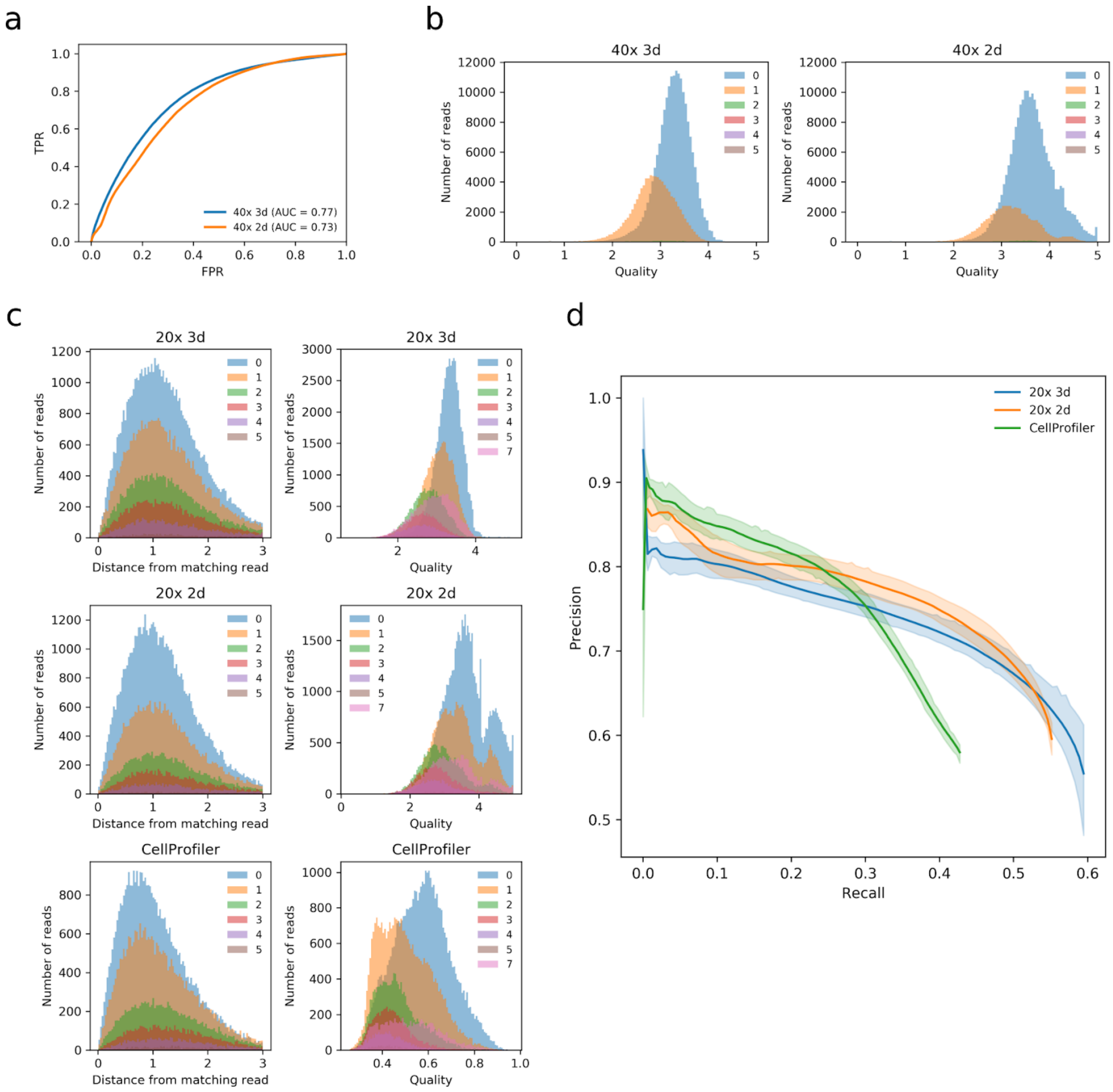


**Supplementary Fig. 10.** Quantitative Evaluation of Image Analysis Pipelines. **a)** Receiver operating characteristic of proposed image decoding pipeline for 3d and 2d analysis of 40x data using list of targeted sequences to evaluate false positive and true positive. **b)** Histograms showing number of reads versus quality for each distribution of decoded reads with number of mismatch respect to a targarged sequence from 0 to the number of sequencing rounds. **c)** Pair of histograms showing for each 20x analysis the distribution of read counts with respect to the distance to the ground truth matching read and read quality. Colors represent number of mismatch with respect to the ground truth matching read (specifically, 0 equal to perfect match, 1 to 5 number of mismatches between the two sequences, 7 represents decoded false positive reads without a matching ground truth read). **d)** Precision versus recall for 20x data image analysis results of 3d and 2d proposed pipeline, and 2d results of CellProfiler pipeline[11]. Precision and recall are evaluated on four equally sized parts of the data at decreasing values of quality threshold, and mean and standard deviation are plotted for each of the three analyses.

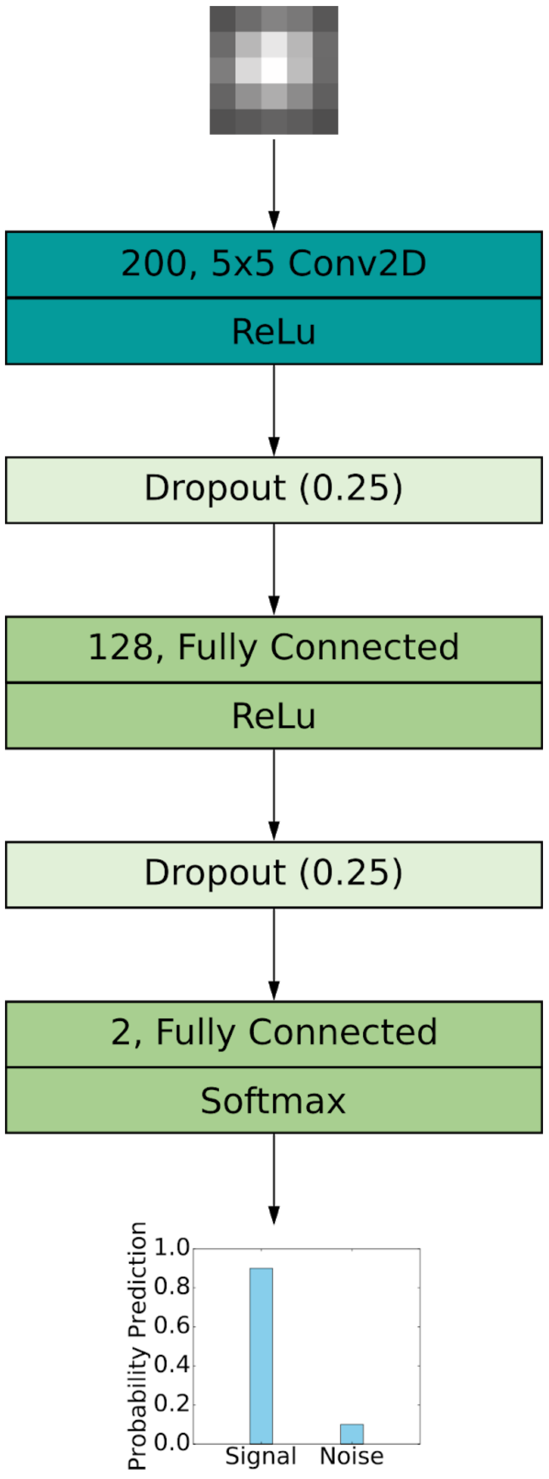


**Supplementary Fig. 11.** Convolutional Neural Network Architecture for Signal Candidate Predictions. The network takes as input a 5x5 pixel window centred in each signal candidate detection and provides probability predictions for the candidate to be signal and noise.

**Supplementary Video.** Video tutorial for interactive inspection and visualization of the decoded spatial gene expression from a mouse brain coronal section through TissUUmaps online viewer (https://tissuumaps.research.it.uu.se/demo/isseq.html).

| | |
|---|---|
| Pattern 0 | Gad1 |
| Pattern 1 | Rgs12 |
| Pattern 2 | Adgrl2, Atp1b1, Kit, Serpini1, Pvalb |
| Pattern 3 | Reln, CdA3 |
| Pattern 4 | Calm2, Rgs4, Pde1a, Nrn1, Nov, Neurod6, Lamp5, Gda, Gabrd, Crym, Crhbp, Cck, Bcl11b, Arpp21, 3110035E14Rik, Vip, Rprm, Nr4a2, Htr3a, Enc1, Cox6a2, Cnr1, Satb1, Snca |
| Pattern 7 | Plcxd2, Sema3c, Sulf2, Slc24a2 |
| Pattern 8 | Aldoc, Ndnf |
| Pattern 11 | Chrm2 |
| Pattern 12 | Calb2, Scg2, Sncg, Tac1, Zcchc12 |
| Pattern 13 | Calb1, Cxcl14, Rorb, Thsd7a, Id2 |
| Pattern 14 | Fos, Grin3a, Nos1, Penk, Rasgrf2, Rgs10, Cpne5, Fxyd6, Gap43, Th, Wfs1 |
| Pattern 15 | Pcp4, Pthlh |
| Pattern 16 | Fam19a1, Pvrl3, Tac2, Cux2, Npy2r |
| Pattern 18 | Enpp2, Plp1, Slc6a1 |
| Pattern 19 | Kcnk2, Synpr |

**Supplementary Table 1a.** Spatial patterns composition of SpatialDE “Automatic Expression Histology” (Supplementary Fig. 6b).

| | |
|---|---|
| Pattern 1 | Fos, Kit, Nrn1, Cck, Neurod6, Satb1, Slc24a2 |
| Pattern 3 | Nos1, Penk, Th, Calb2, Fxyd6, Gap43, Sncg, Zcchc12 |
| Pattern 4 | Rprm, 3110035E14Rik, Bcl11b, Crym, Enc1, Gda, Nov, Pde1a |
| Pattern 5 | Scg2, Tac1 |
| Pattern 6 | Enpp2, Plp1 |
| Pattern 9 | CdA3, Npy2r, Synpr, Calb1, Cox6a2, Wfs1 |
| Pattern 10 | Tac2 |
| Pattern 11 | Calm2, Fam19a1, Arpp21, Cux2, Gabrd, Lamp5, Rgs4, Vip |
| Pattern 12 | Id2 |
| Pattern 14 | Cnr1, Crhbp, Htr3a, Nr4a2, Snca, Cxcl14, Rgs12 |
| Pattern 15 | Adgrl2, Atp1b1, Serpini1, Sulf2, Chrm2, Gad1, Pcp4, Pvalb |
| Pattern 18 | Aldoc, Cpne5, Grin3a, Kcnk2, Pthlh, Rasgrf2, Reln, Rgs10, Sema3c, Slc6a1, Thsd7a, Plcxd2, Pvrl3, Rorb |
| Pattern 19 | Ndnf |

**Supplementary Table 1b.** Spatial patterns composition of SpatialDE "Automatic Expression Histology" (Supplementary Fig. 6c).
